# Increases in Arctic extreme climatic events are linked to negative fitness effects on the local biota

**DOI:** 10.1101/2024.09.26.615176

**Authors:** M Lemaire, S. Bokhorst, A. Witheford, M. Macias-Fauria, R. Salguero-Gomez

## Abstract

The Arctic harbours uniquely adapted biodiversity and plays an important role in climate regulation. Strong warming trends in the terrestrial Arctic have been linked to an increase in aboveground biomass (Arctic greening) and community-wide shifts such as the northwards-expansion of boreal species (borealisation). Whilst considerable efforts have been made to understand the effects of warming trends in average temperatures on Arctic biota, far fewer studies have focused on trends in extreme climate events and their biotic effects, which have been suggested to be particularly impactful during the Arctic winter months. Here, we present an analysis of trends in two ecologically-relevant winter extreme events –extreme winter warming and rain-on-snow, followed by a meta-analysis on the evidence base for their effects on Arctic biota. We show a strong increase in extreme winter warming across the entire Arctic and high variability in rain-on-snow trends, with some regions recently experiencing rain-on-snow for the first time whilst others seeing a decrease in these events. Ultimately, both extreme events show significant changes in their characteristics and patterns of emergence. Our meta-analysis –encompassing 178 effect sizes across 17 studies and 49 species– demonstrates that extreme winter warming and rain-on-snow induce negative impacts on Arctic biota, with certain taxonomic groups –notably angiosperms and chordates (mostly vertebrates)– exhibiting higher sensitivity than others. Our study provides evidence for both emerging trends in Arctic winter extreme climate events and significant negative biotic effects of such events –which calls for attention to winter weather variability under climate change in the conservation of Arctic biodiversity, whilst highlighting important knowledge gaps.

## 2. Introduction

Extreme climate events are characterised by climatic conditions that are rare for a given location and time. A climatic event is classified as ‘extreme’ when it falls at the tail end of the frequency distribution for a given observed variable, normally defined by a percentile (e.g., 1^st^, 5^th^, 10^th^, 90^th^, 95^th^, 99^th^; National Academies Press, 2016). Gradual warming trends have been linked to increased frequency and intensity of extreme climate events: over 400 studies have linked rises in extreme climate events to climate change, especially temperature increases (CarbonBrief, 2022). As gradual warming trends increase the intensity and frequency of extreme events, the frequency with which the physiological tolerance boundary for organisms is crossed is expected to increase (Harris *et al.,* 2018). Since organisms have become adapted to local climatic variation through temporal scales much longer than the ones under which current climatic extremes are emerging (Gregory, 2009), extreme climate events pose tangible risks to their viability (Harris *et al.,* 2018). Ultimately, the crossing of tolerance boundaries may result in population declines, migration-led range shifts, and local or total extinctions (Smale & Wernberg, 2013; Boucek & Rehage, 2014).

The Arctic tundra is the terrestrial biome experiencing the greatest warming rates resulting from climate change (Rantanen *et al*., 2022). Arctic warming has been reported to deviate between twice (Walsh, 2014) to four times from the global average temperature (Rantanen *et al*., 2022). These strong trends have been linked to increasing aboveground biomass in the tundra, known as “Arctic greening” (Myneni *et al*., 1997; Berner *et al.,* 2020; Myers-Smith *et al*., 2020) and to community shifts as boreal species expand northwards (Fossheim *et al*., 2015; Speed *et al*., 2021). Post and colleagues (2009) argued that the perception of low species richness in the Arctic has led to paucity in biodiversity-related research efforts in this area. However, this perception fails to recognise the unique adaptations of Arctic biota and the low functional redundancy of species in Arctic ecosystems, making each species crucial as functional redundancy ensures ecological resilience (Biggs *et al*., 2020).

Most studies examining the effects of rapid Arctic climate change have thus far focused on growing season fitness, with responses to winter warming being under-researched and relatively unknown, despite their biological and ecological importance (Bjerke, 2009). In terrestrial systems, Arctic biota has adapted to extreme cold by slowing down or stopping basic physiological processes when temperatures drop in the autumn. For instance, plants, mosses, and lichens adjust their lipid composition in the dormant season to prevent denaturation of their lipid membranes by freezing temperatures (Dalmannsdóttir *et al*., 2001; Chen *et al*., 2013; Strimbeck *et al*., 2015). These adaptations respond to environmental cues, with lipid membrane composition reverting in warmer months. Such adaptations suggest that Arctic biota might be very sensitive to short-lived extreme climatic events in the cold season, since it is unclear whether Arctic species can re-acclimate to winter temperatures following short winter warming events in which de-acclimation is triggered (Bokhorst *et al*., 2023). For example, plant species may exhibit advanced phenology in response to extended warming events (Ladwig *et al*., 2019) with early emergence from dormancy during winter likely being a key driver in vascular plant mortality (Bokhorst *et al*., 2010). Ultimately, differences in life cycles, growth forms, and adaptive strategies may underlie why some species and taxonomic groups are more resistant to extreme winter warming than others (Bjerke *et al*., 2011).

Climate change is increasing the number of extreme events in the Arctic, in particular during the winter season (Vikhamar-Schuler et al., 2016). Arctic warming may increase the frequency of extreme climatic events such as rain-on-snow and extreme winter warming. Rain-on-snow events occur when rain falls on existing snowpacks before refreezing, potentially forming hard crusts of ice (Serreze *et al.,* 2021). Both rain-on-snow and extreme winter warming may lead to the formation of ice layers in the snowpack with key consequences for ecosystem functioning (Hansen *et al*., 2013; Bartsch *et al*., 2023). Rain-on-snow has been shown to accentuate food shortages for large herbivores in Arctic regions, such as in Siberia and Svalbard, leading to large-scale die-offs (Forbes *et al*., 2016) and trophic cascade collapses (Stien *et al*., 2012; Layton-Matthews *et al*., 2023). Extreme winter warming events rapidly melt snow, exposing the ecosystem to unseasonably warm air. Once the extreme heat ends, Arctic vegetation is subjected to much lower temperatures than they would generally experience as the insulating snow cover is lost. Extreme winter warming events have been linked to higher shoot mortality (Bokhorst *et al.,* 2011), decreased flower abundance (Semenchuk *et al.,* 2013), and reduced reproductive success (Bokhorst *et al*., 2011).

Although the rise in the occurrence of rain-on-snow and extreme winter warming events has been reported (Forbes *et al*., 2016, Pascual & Johansson, 2022; Bartsch *et al*., 2023), a systematic, Arctic-wide quantification of said increases is still lacking. The knowledge gap on the overall direction, strength, and specificity of the biotic effects of extreme winter climate events makes research into both winter warming and rain-on-snow of vital importance. Synthesising our current understanding of the impacts of extreme winter events on Arctic biota is thus key to determining their overall impact – and threats – on Arctic terrestrial ecosystems’ stability. Here, we conduct a two-part integrative analysis to address three aims. First, we analyse atmospheric hourly reanalysis climate data (ERA5) for the terrestrial Arctic (Hersbach *et al.,* 2023), using data from the European Centre for Medium-Range Weather Forecasts (ECMWF) to (1) determine whether the frequency and intensity of extreme winter events in the Arctic have increased since 1950. Second, we carry out a meta-analysis to (2) investigate whether rain-on-snow and extreme winter warming events in the Arctic are negatively correlated with the fitness of native biota. We then use a meta-regression to (3) assess the impacts of methodological and biological moderators in modulating the reported effects of extreme winter events on the fitness of Arctic biota. Through this integrated approach, we aim to enhance our understanding of the complex interactions between climate change, extreme events, and Arctic biota, whilst also highlighting important knowledge and research gaps.

## 3. Methods

### 3.1 Overview

To assess the current impacts of climate change in the Arctic on extreme winter climate events and terrestrial biota, we conducted a two-part study. We first determined whether the frequency and intensity of extreme climate events in Arctic tundra have shown a significant increase since the baseline period 1950-1980 using ECMWF ERA5 data (Hersbach *et al.,* 2023). Then, we examined whether extreme winter warming and rain-on-snow events have a significant effect on the fitness of Arctic biota by conducting a meta-analysis (Gurevitch *et al.,* 2018; O’dea *et al.,* 2021). By linking ecological responses to meteorological data, we provide a comprehensive understanding of the impacts of climate change on the Arctic.

### 3.2 Climate Analysis: Frequency and Intensity of Extreme Climate Events

To analyse trends in extreme winter warming events, we used ERA5 hourly data (Hersbach *et al.,* 2023). We defined the winter season for the Arctic as October-March (Rennert *et al*., 2009) and used the geographical boundaries for the Arctic region established by Martin *et al*., (2022) (Appendix 1). We aggregated ERA5 data to daily intervals and 1° latitude by 1° longitude grid cells before our analysis. To identify extreme winter warming days, we calculated the 99^th^ percentile for the maximum daily winter temperature for each grid cell in the Arctic based on the climatological period 1950-1980. We classified winter days in which the maximum temperature exceeded this cut-off as an extreme winter warming event. We calculated annual cumulative warming exceedance by adding the daily differences between the maximum temperature of an extreme winter warming event and the threshold for extreme winter warming for all extreme winter warming events in a grid cell over a year. A single extreme event was defined as a group of consecutive winter warming days within the same 1° by 1° grid cell.

We next analysed trends in extreme winter warming events and identified vulnerable regions. To establish whether there was a temporal trend in extreme winter warming events, we performed a non-parametric Theil-Sen regression (Sen, 1968) on event length, intensity, maximum temperature, and annual event number. Here, intensity refers to the cumulative warming exceedance and is calculated by multiplying the exceedance value of each day by its corresponding day number within an event. For instance, the intensity of a two-day event that experiences maximum daily temperatures of 5 and 7°C, with an extreme warming threshold set at 3 °C is ((5 − 3) × 1) + ((7 − 3) × 2) = 10. To identify which areas experienced most increases in extreme winter warming events, we plotted the mean annual cumulative winter warming exceedance per grid cell for 1950-1980 and 1990-2020. To determine statistically significant differences between both periods, we performed paired t-tests at pan-Arctic and regional scales, after a Mantel test on our grid data demonstrated positive but weak spatial autocorrelation (r = 0.077, *p* < 0.010).

We then analysed trends in rain-on-snow events and identified vulnerable regions. Here, we defined rain-on-snow events as 3 mm of rain in a day with a mean snow depth of 3 mm of snow water-equivalent, with maximum daily temperature greater than 1°C in the Arctic winter season (adapted from Rennert *et al*., 2009). To do so, we used daily total rainfall, mean daily snow depth, and maximum daily temperature from the ERA5 dataset (Hersbach *et al.,* 2023), previously aggregated to 1°×1° resolution. To determine whether each grid point was independent, we performed a Mantel test, which produced a significant but weak signal (r = 0.090, *p* < 0.010). To examine temporal trends in total rain-on-snow, event length, and intensity, we again used Theil-Sen regression. In this case, we measured intensity as the sum of total daily rainfall on an event multiplied by the day of the event. For example, the intensity of a two-day rain-on-snow event with 5 and 7 mm of rain on subsequent days is (5 × 1) + (7 × 2) = 19. We identified regional patterns of rain-on-snow occurrence across the Arctic by plotting mean total annual rain-on-snow (mm) for each grid cell for the periods 1950-1980 and 1990-2020. Finally, we performed a paired t-test to determine the statistical significance of difference between both periods, both at pan-Arctic and regional scales.

### 3.3 Meta-analysis

#### Search Protocol

To determine whether extreme winter events in the Arctic tundra have a significant effect on its biota, we conducted a meta-analysis. To ensure the reproducibility of our meta-analysis, we used a systematic search protocol outlined in the PRISMA-EcoEvo guidelines (O’Dea *et al*., 2021) on SCOPUS and Web of Science. First, to capture all the relevant literature, we developed a comprehensive search string using the most commonly occurring keywords in relevant papers via a scoping search. We performed the scoping search in Google Scholar with the keywords “extreme climate event*” and “Arctic tundra”. To identify the most commonly occurring keywords, we selected the first 10 relevant papers from the scoping search and produced a word frequency list in descending order of frequency (Appendix 2). We included these keywords and their synonyms in the search string. Further refinement, as detailed in Appendix 3, then took place to reduce the number of papers not relevant to the research question.

The literature search was conducted in SCOPUS and Web of Science on the 15^th^ of November 2023 and the 11^th^ of March 2024, respectively. Using the aforementioned search strings, we obtained 226 hits from SCOPUS and 154 from Web of Science. These hits produced 276 unique papers after deduplication. We screened the hits using pre-determined inclusion criteria, developed so that only pertinent studies would be included in our meta-analysis. Namely, the study had to (1) be an original research article so our meta-analysis would synthesise the current state of the literature; (2) quantify the response of a fitness-related trait (*e.g*., body mass, photosynthetic rate) in a native species to either extreme winter warming or rain-on-snow; (3) have been conducted within the Arctic region, following the spatial definition by Martin *et al*. (2022) (Appendix 1); and (4) report sample sizes, means, and standard deviations for both ‘control’ and ‘response’ groups. In experimental studies with multiple treatments (*e.g*., canopy warming, and canopy + soil warming), only the most extreme treatment was extracted to avoid pseudo-replication. We only used studies published in English so that we could extract the data without needing professional translation.

#### Data Collection

We employed two additional methods to extract the relevant data from studies where means and standard deviations of fitness-related responses to rain-on-snow or winter warming were not explicitly reported. First, for studies where the data were available in figures, we used ImageJ (Schneider *et al*., 2012) for data extraction. Second, when not available in the publication, including on figures, we requested the data from the authors. In cases where these methods were unsuccessful (19%), we excluded the papers from our meta-analysis (n = 4 out of 21).

To derive means and standard deviations from observational studies measuring a fitness-related variable over time, data were categorised into ‘response’ (extreme) and ‘control’ groups. If a study explicitly mentioned which years experienced extreme events, these were assigned to the ‘response’ category, with the years preceding an extreme year classed as ‘control’. If extreme years were not highlighted in the publication, we established a threshold based on the environmental variable recorded in the study. To do so, we calculated the 99^th^ percentile based on the climatological period 1950-1980 in a 5° × 5° region around the study location from ERA5 data (Hersbach *et al.,* 2023). Extreme years exceeding this threshold were only selected if a ‘control’ year data point was available for the year preceding the extreme year. The mean and standard deviations were then calculated for each category, with the sample size representing the number of years in each.

We identified 17 studies (Appendix 4) with relevant data on the effects of extreme winter events on fitness-related traits in Arctic biota. An independent assessor performed abstract and full-text screening to assess reproducibility in our pipeline. To determine the inter-rate agreement, we used Cohen’s Kappa statistic, which indicates the level of agreement between two raters beyond the level which would be expected by chance (Koricheva *et al.,* 2013) (Cohen’s Kappa= 0.88).

Moderators are factors that could potentially influence the direction and strength of the effect sizes being analysed. To enable comparisons between different moderators across studies, for each study we systematically recorded (1) whether the study examined the effects of rain-on-snow or extreme-winter-warming, (2) whether it was an experimental or observational study, and (3) the year and location of the study. We also documented the species name, taxonomic group, phylum, and response variable name (*e.g.,* shoot growth (Treharne *et al.,* 2018), photosystem II – PSII activity (Bokhorst *et al.,* 2023). To include response as a valuable moderator, meaning a variable which may explain variability between studies, (*sensu* Gu *et al*., 2023), we divided response variables into two broader categories: quantity and energetic requirements (Appendix 5). Quantity-related response variables capture the impacts of changes in life-history traits on populations and individuals (*e.g.,* abundance (Bokhorst *et al.,* 2012), calf at heel (Loe *et al.,* 2016)). Following extreme events, changes in size, abundance, or density may occur as a result of changing reproductive rates, migration, and/or mortality (Harris *et al.,* 2018). Quantifying these changes helps assessing the population-level impacts and how these may shape ecosystem dynamics. Energy-related responses variables encompass variables linked to physiological processes and metabolic demands that adjust in response to environmental changes (*e.g.,* photosynthetic rate (Bokhorst *et al.,* 2018), time required to dig to layer C (Poirier *et al.,* 2021)). Extreme events may impose significant energetic costs on individuals, leading to changes in metabolic rates and resource allocation.

We assessed the effects of methodological moderators to determine whether different studies could be analysed together and test for potential biases. As a single author (Bokhorst) was present in nine of the seventeen studies included (Bokhorst *et al*., 2008; 2010; 2011; 2012a; 2012b; 2015; 2018; Appendix 4), we determined whether there was a significant difference in reported effect sizes between studies authored by Bokhorst *et al*. and the rest. This assessment was done to ensure that no bias was being introduced through the uneven distribution of first authorship. To assess the impact of study type on effect size, we compared different methodological approaches.

#### Data Analysis

To facilitate the synthesis of findings across multiple studies, we calculated effect sizes for each study using the standardised mean difference with heteroscedastic population variances (SMDH). This statistical method, also known as Hedge’s *d* (Hedges, 1981), considers the difference in variance between the two compared groups and is widely used in meta-analyses (Bonett, 2009). Effect sizes and corresponding sampling variances were calculated using the ‘*escalc*’ function in the *metafor* R package (Viechtbauer, 2010). From the 17 examined studies, we calculated 183 effect sizes across 49 species. To ensure consistency in the directionality of effect sizes, effect sizes were multiplied by -1 when an increase in response mean affects fitness negatively (*e.g.,* winter mortality (Loe et al, 2016)). Outliers were identified and removed using the ‘metaoutliers’ function from the ‘altmeta’ package in R (Lin *et al.,* 2022), which classifies an effect size as an outlier if, in terms of absolute magnitude, its standardised residual is greater than three. Following the exclusion of outliers, we were left with 178 effect sizes across 17 studies and 49 species (Supplementary Material 2).

To calculate and test for an overall effect size, we used a meta-analytical (‘null’) model with the effect size (SMDH) as the response variable. A ‘null’ model does not include any moderators and examines the overall effect size across all studies, providing a general estimate without accounting for the effects of specific moderators. In this step, we used the ‘*rma.mv*’ function from the *metafor* R package. To control for non-independence of effect sizes, we included study ID, effect size ID, species name, and a correlation matrix quantifying phylogenetic relatedness as random effects in the null model (Noble *et al*., 2017; Cinar *et al.,* 2022). We used data from the OpenTreeOfLife (Hinchliff *et al*., 2015) with the *ape* (Paradis & Schliep, 2019) and *rotl* R packages (Michonneau *et al.,* 2016) to build a phylogenetic tree and produce the correlation matrix.

To test the effects of various moderators on the impacts of extreme events on fitness-related responses of the Arctic biota, we used meta-regression analysis using the ‘*rma.mv*’ function in the *metafor* R package. In a meta-regression, the effect sizes are regressed against one or multiple moderator variables, thus allowing for the quantification of the association between moderators and effect sizes. For each model, we used the ‘*r2_ml*’ function from the *orchaRd* R package (Nakagawa *et al*., 2023) to obtain the total heterogeneity (Q_M_) and the proportion of this that is explained by the moderators (marginal R^2^) (Bates *et al*., 2014). All meta-regression models included the same random effects as in the null model and SMDH as the response variable. To test whether each moderator showed evidence for differing effects of extreme events on Arctic biota, we created models without an intercept.

We conducted all statistical analyses in R v 4.3.2 (R core Team, 2023) and have made them available online: https://github.com/mayalemaire/Arctic_Climate_Project

#### Areas of special interest

We identified regions of interest during the abstract screening stage of the literature review process. Whilst our climatic analysis spans the whole Arctic, we present all analyses with a focus on the four regions most prevalent in the literature on biotic impacts of winter extreme events, since they informed our meta-analysis. These are Northern Alaska, Northern Fennoscandia, Svalbard, and Yamal (North-western Siberia).

## 4. Results

### 4.1 Climate Analysis

#### Extreme Winter Warming

Over the period 1990-2020, annual cumulative winter warming exceedance has been greater than that over the baseline period spanning 1950-1980, with a mean increase of 2.8 °C in winter warming exceedance (*p* < 0.001, Fig 1). We found a statistically significant increase in annual cumulative winter warming between the two periods across all four areas of special interest (Northern Alaska: mean increase = 2.9 °C, *p* < 0.001; Northern Fennoscandia: mean increase = 3.0 °C, *p* < 0.001; Svalbard: mean increase = 6.6 °C, *p* < 0.001; Yamal: mean increase = 7.2 °C, *p* < 0.001; Fig 1).

**Fig. 1.**
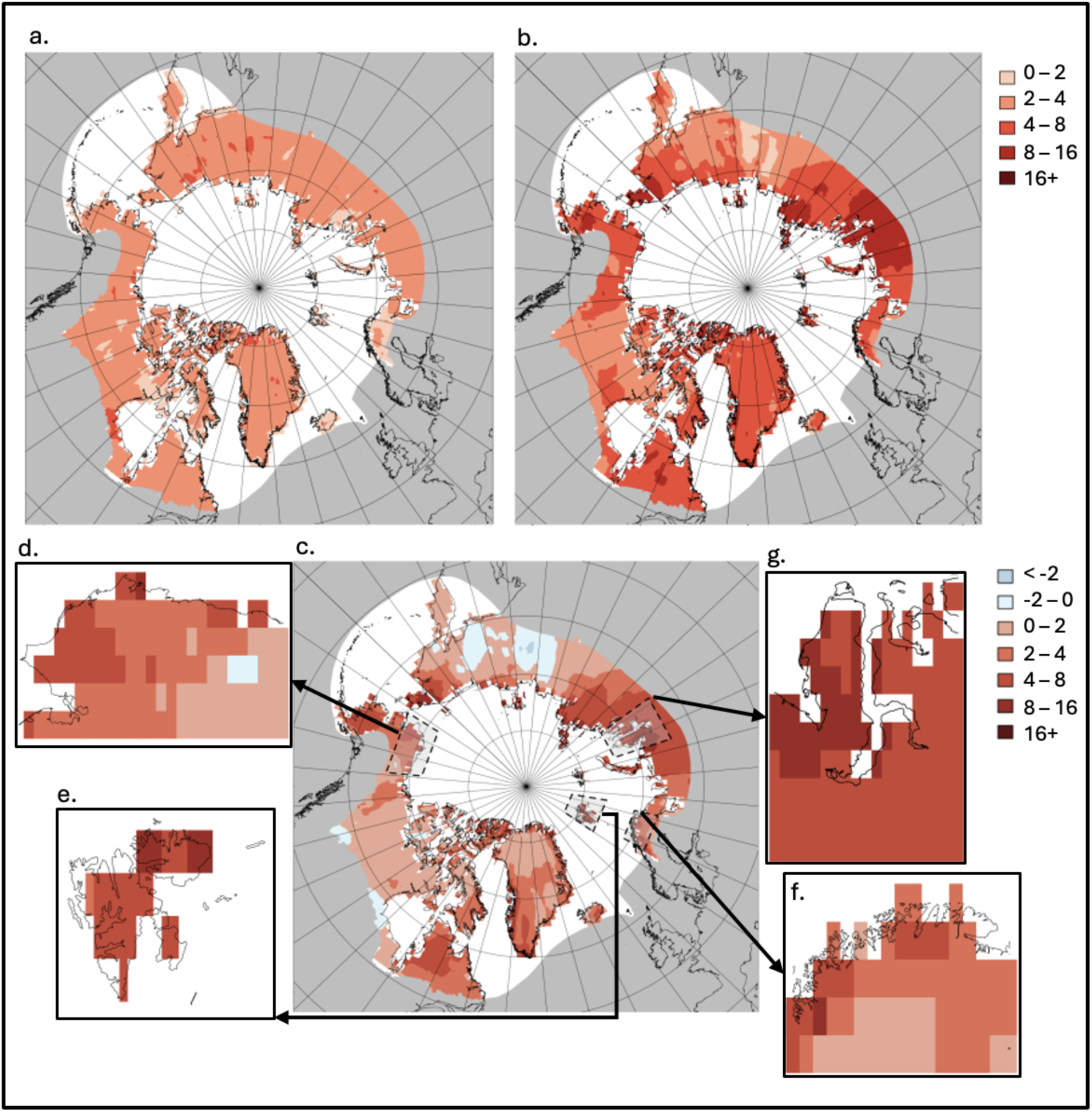
The Arctic has experienced a significant increase in annual cumulative winter warming exceedance. Average cumulative winter –defined as October-March, see *Methods*– warming exceedance (°C) for the (**a**) baseline period 1950-1980 and (**b**) modern period 1990-2020. The differences between the two periods (°C) are shown in (**c**). The grey boxes in (**c**) indicate areas of special interest, expanded in: (**d**) Northern Alaska, (**e**) Svalbard, (**f**) Northern Fennoscandia, and (**g**) North-western Siberia (Yamal). Spatial autocorrelation was assessed using a Mantel test (r = 0.077, *p* < 0.010).

The increase in winter warning exceedance is in line with measured gradual warming trends across the Arctic (Appendix 6). In the analysis of gradual warming trends in winter temperatures, a statistically significant increase in mean daily maximum temperature during winter seasons was found in all regions. Svalbard experienced the most pronounced gradual warming trend, with a mean increase of 0.12 °C in average maximum daily temperature for each winter season (i.e., per year. *p* < 0.001). This was followed by Northern Alaska and Yamal, which both experienced an increase of 0.06 °C in maximum daily winter temperature per winter season (*p* < 0.001). Northern Fennoscandia experienced the weakest trend, with 0.03 °C per winter season (*p* < 0.001).

Extreme winter warming events have become longer, hotter, and more intense every year in all areas of special interest (Fig. 2). The area experiencing the greatest positive trend in extreme warming event length, intensity, and maximum temperature reached is Yamal. In this area, there was a positive increase in extreme event length of 0.029 days per winter season (i.e., per year, *p* < 0.001) since 1950. Extreme winter warming event intensity in Yamal increased at a rate of 0.117 per season (*p* < 0.001) and maximum temperature reached in warming event showing an increase of 0.016 °C per season (*p* < 0.010). In all regions, there was a significant increase in event length (Northern Alaska: mean increase = 0.025 days per season, *p* < 0.001; Northern Fennoscandia: mean increase = 0.006 days per season, *p* < 0.001; Svalbard: mean increase = 0.015 days per season, *p* < 0.001). Extreme winter warming intensity increased in all regions (Northern Alaska: mean increase = 0.091 per season, *p* < 0.001; Northern Fennoscandia: mean increase = 0.029 per season, *p* < 0.001; Svalbard: mean increase = 0.032 per season, *p* < 0.001). Maximum temperature reached in an extreme winter event showed a significant increase in all regions but Svalbard (Northern Alaska: mean increase = 0.004 °C per season, *p* < 0.05; Northern Fennoscandia: mean increase = 0.007 °C per season, *p* < 0.001).

**Fig. 2.**
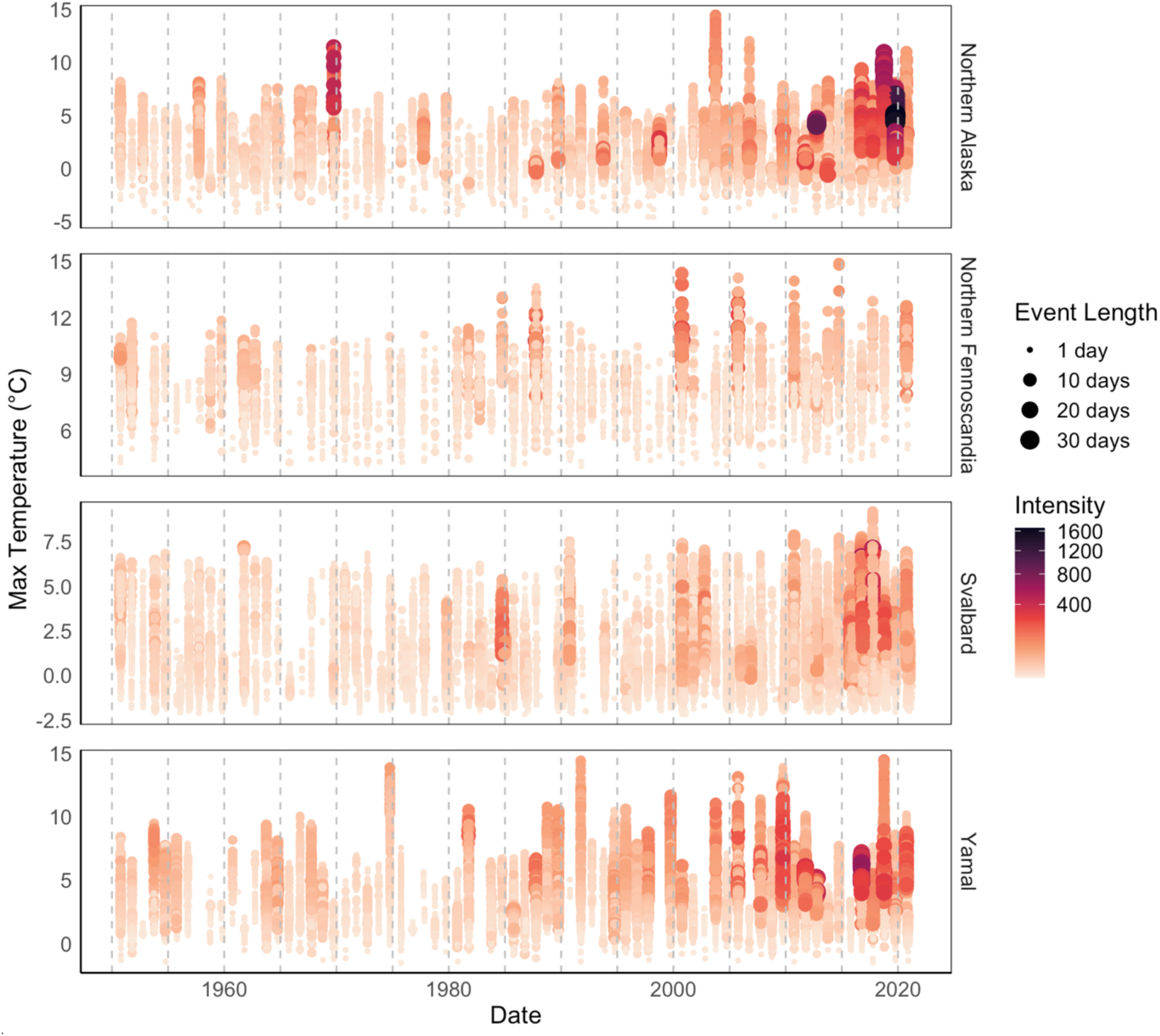
Yearly extreme winter warming events since 1950. The plot depicts individual extreme winter warming events detected on a grid-point basis in different Arctic areas of special interest. Maximum temperature (°C) reached during each event is represented on the y-axis, with colour indicating intensity and size representing the length of the event. Regions are ordered alphabetically from top to bottom: Northern Alaska, Northern Fennoscandia, Svalbard, and Yamal (North-western Siberia). In all regions apart from Northern Fennoscandia, the intensity and length of extreme winter warming events showed a statistically significant increase.

#### Rain-On-Snow

At the pan-Arctic level, there was no significant increase in total yearly rain-on-snow in the period 1990-2020, compared to the baseline period of 1950-1980 (mean difference = 0.130 mm, *p* = 0.143) (Fig. 3). However, while there have been localised regions with decreases in rain-on-snow, mostly in warm and wet southern Arctic locations and Eastern Siberia, many Arctic regions have experienced an increase in rain-on-snow, while most of the Arctic shows no trend, driving the statistical non-significance. Eleven percent of all the grid points across the Arctic where at least one rain-on-snow event was detected in the period 1990-2020 did not experience any in the baseline period 1950-1980. On the other hand, 9.8% of grid points in which rain-on-snow was identified in the baseline period did not have any events in the modern period. In the areas of special interest, Northern Alaska, Svalbard, and Yamal experienced a significant increase in mean yearly rain-on-snow (Alaska: mean difference = 0.384 mm, *p* < 0.010; Svalbard: mean difference = 2.338, *p* < 0.005; Yamal: mean difference = 0.172 mm, *p* < 0.005), whereas yearly rain-on-snow decreased in Northern Fennoscandia between the two periods (mean difference = −2.546 mm, *p* < 0.010).

**Fig. 3.**
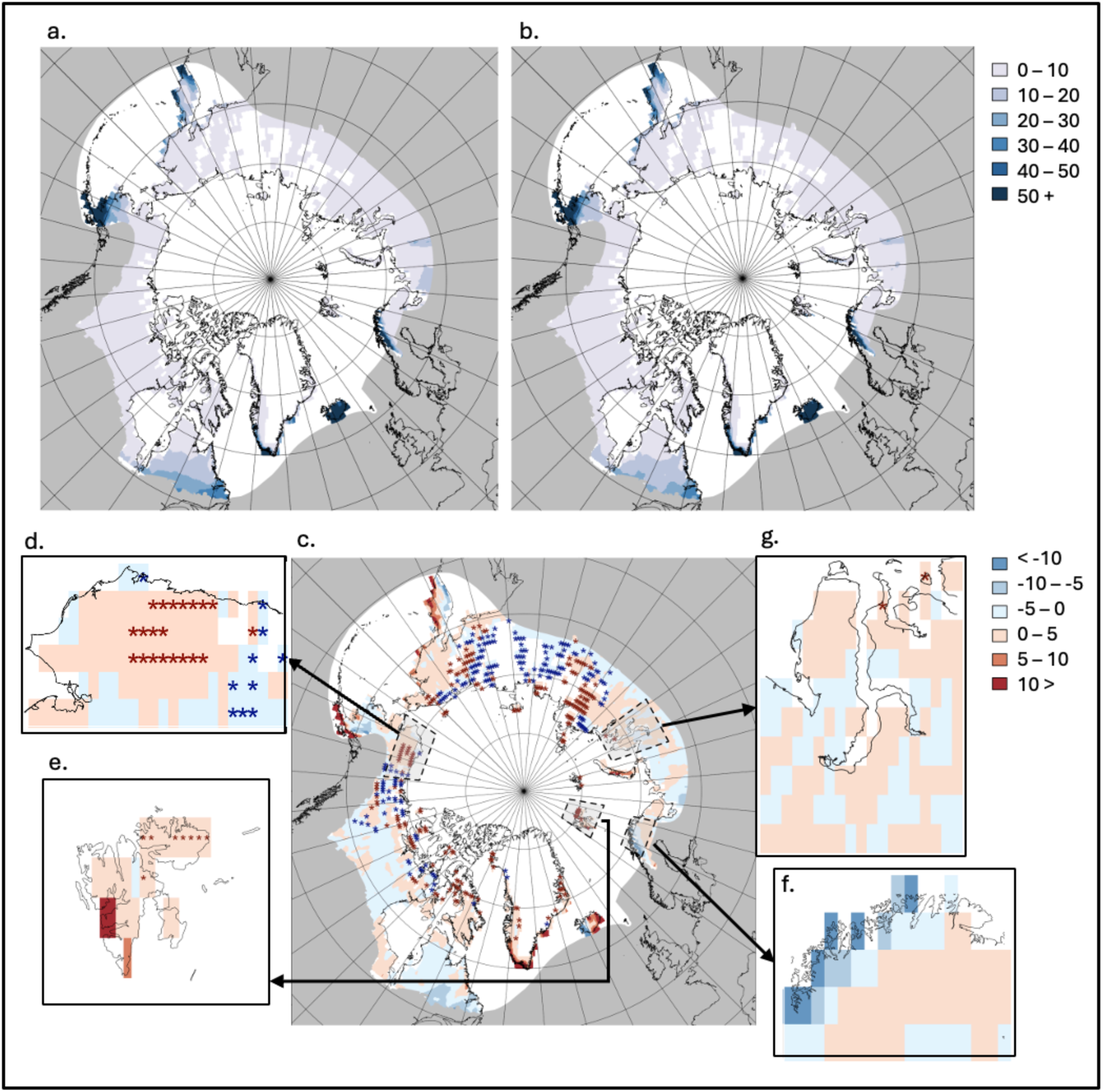
The change in mean total rain-on-snow events across the Arctic. Average total annual rain-on-snow (mm) for the (**a**) baseline period 1950-1980 and (**b**) modern period 1990-2020. The differences between the two periods (mm) are shown in (**c**). Red stars indicate grid points where rain-on-snow events have been detected in the modern period but not in the baseline period whilst blue stars indicate the opposite. The grey boxes in (**c**) indicate areas of special interest, expanded in: (**d**) Northern Alaska, (**e**) Svalbard, (**f**) Northern Fennoscandia, and (**g**) Yamal (North-western Siberia). Spatial autocorrelation was assessed using a Mantel test (r = 0.090, *p* < 0.010).

There were significant positive trends in yearly precipitation across the different Arctic regions of interest (Appendix 7). There are significant positive linear trends in average total winter rainfall over time for all regions (Northern Alaska: slope = 0.039 mm per season, *p* < 0.001; Northern Fennoscandia: slope = 0.466 mm per season, *p* < 0.001; Svalbard = 0.235 mm per season, *p* < 0.001; Yamal: slope = 0.168 mm per season, *p* < 0.001). There was a similar trend for average total winter snowfall over time (Northern Alaska: slope = 0.298 mm per season, *p* < 0.001; Northern Fennoscandia: slope = 0.290 mm per season, *p* < 0.001; Svalbard = 1.087 mm per season, *p* < 0.001; Yamal: slope = 0.391 mm per season, *p* < 0.001). We found statistically significant positive trends in the proportion of rain to snow over winter seasons in all regions (Northern Alaska: slope = 0.0003 rain:snow per season, *p* < 0.001; Northern Fennoscandia: slope = 0.002 rain:snow per season, *p* < 0.001; Svalbard = 0.001 mm per season, *p* < 0.001; Yamal: slope = 0.001 mm per season, *p* < 0.001).

Overall, we found no dominant trends in rain-on-snow across Arctic areas of special interest (Fig. 4). However, this is due to the high spatial heterogeneity in the statistically significant individual trends for several regions and across different aspects of rain-on-snow events. In Svalbard, we found no significant trends in average rain-on-snow event length, intensity, or total rainfall, but we did find a significant positive trend in event frequency, with the number of events increasing at a rate of 0.351 events per season (*p* < 0.001). In Northern Fennoscandia and Yamal, we found significant negative trends in rain-on-snow event length, intensity, and total rainfall (Northern Fennoscandia: length = −0.002 days per season, *p* < 0.005; intensity = - 0.00006 per season, *p* < 0.001; total rainfall = −0.021 mm per season, *p* < 0.001; Yamal: length = −0.0001 days per season, *p* < 0.001; intensity = -9.498e-06 per season, *p* < 0.050; total rainfall = −0.007 mm per season, *p* < 0.050). In Northern Fennoscandia, these trends appear driven coastal regions to the west (Fig. 3f). In Northern Alaska, we found a statistically significant decrease in rain-on-snow event total rainfall (slope = -0.0186 mm per season, *p* < 0.05).

**Fig. 4.**
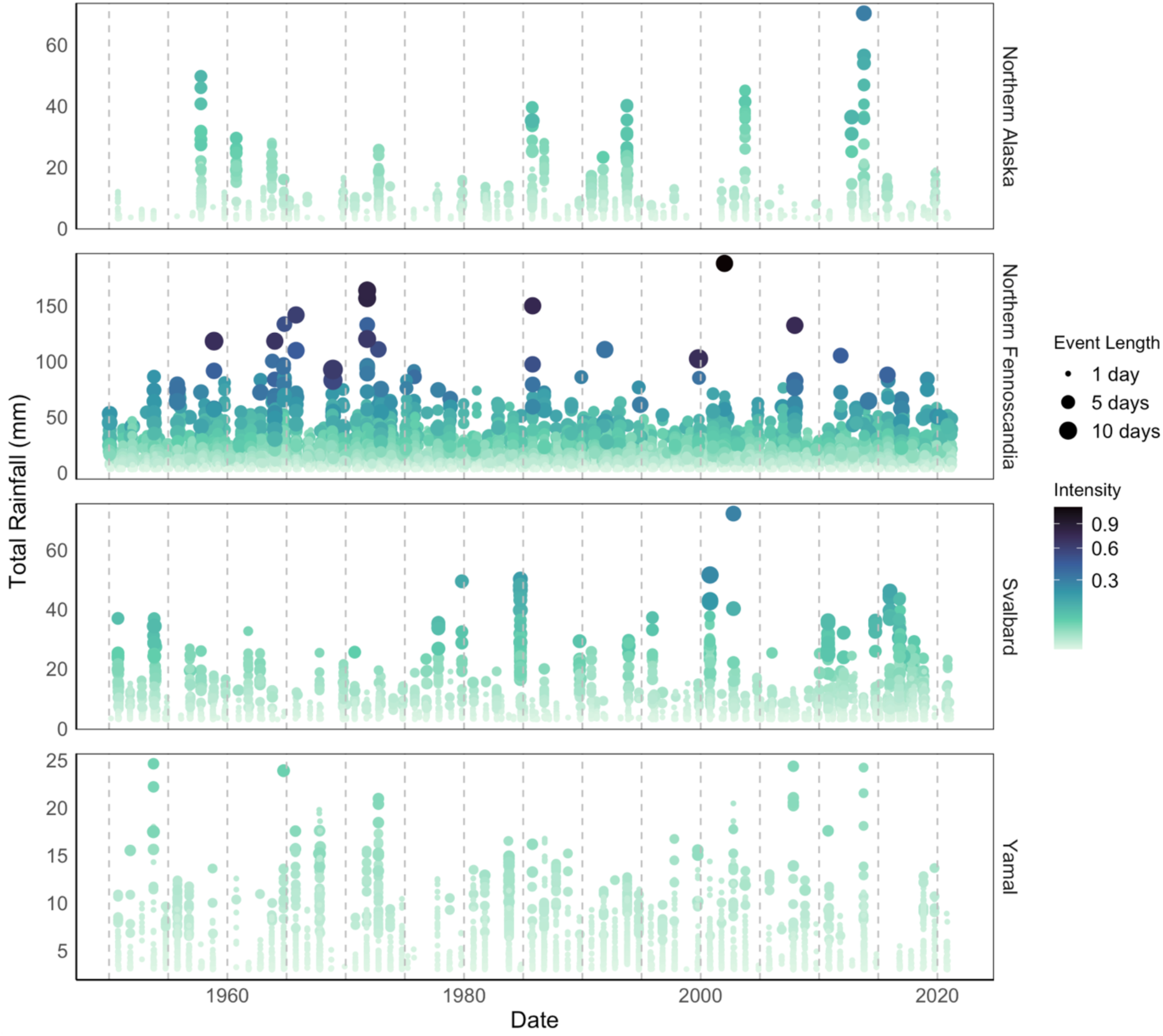
Yearly rain-on-snow events across four Arctic regions since 1950. The plot depicts individual rain-on-snow events detected on a grid-point basis in the different Arctic areas of special interest. Total rainfall (mm) for each event is represented on the y-axis, with colour indicating intensity and size representing length of the event. Regions are ordered alphabetically from top to bottom: Northern Alaska, Northern Fennoscandia, Svalbard, and Yamal (North-western Siberia).

### 4.2 Meta-analysis

Our meta-analysis ‘null’ model (no moderators) revealed that the performance of Arctic biota is negatively impacted by extreme winter events (mean: −0.883, *p* < 0.001, Fig. 5). However, the dataset shows high heterogeneity (I^2^= 85.9%), with 69.1% of effect size variation being attributed to differences between effect sizes, 13.7% to differences between studies, 3.1% to phylogenetic relatedness, and no effect size variation (0.0%) linked to between-species differences. The full model (including all moderators) failed to indicate a significant effect of extreme winter events on Arctic biota. It also fails to explain the heterogeneity in the null model (marginal R^2^ = 6.7%, Q_Mdf=9_ = 3.4, *p* = 0.948). This indicates that none of the moderators have a significant impact on the evidence for the impact of extreme events on Arctic biota.

**Fig. 5.**
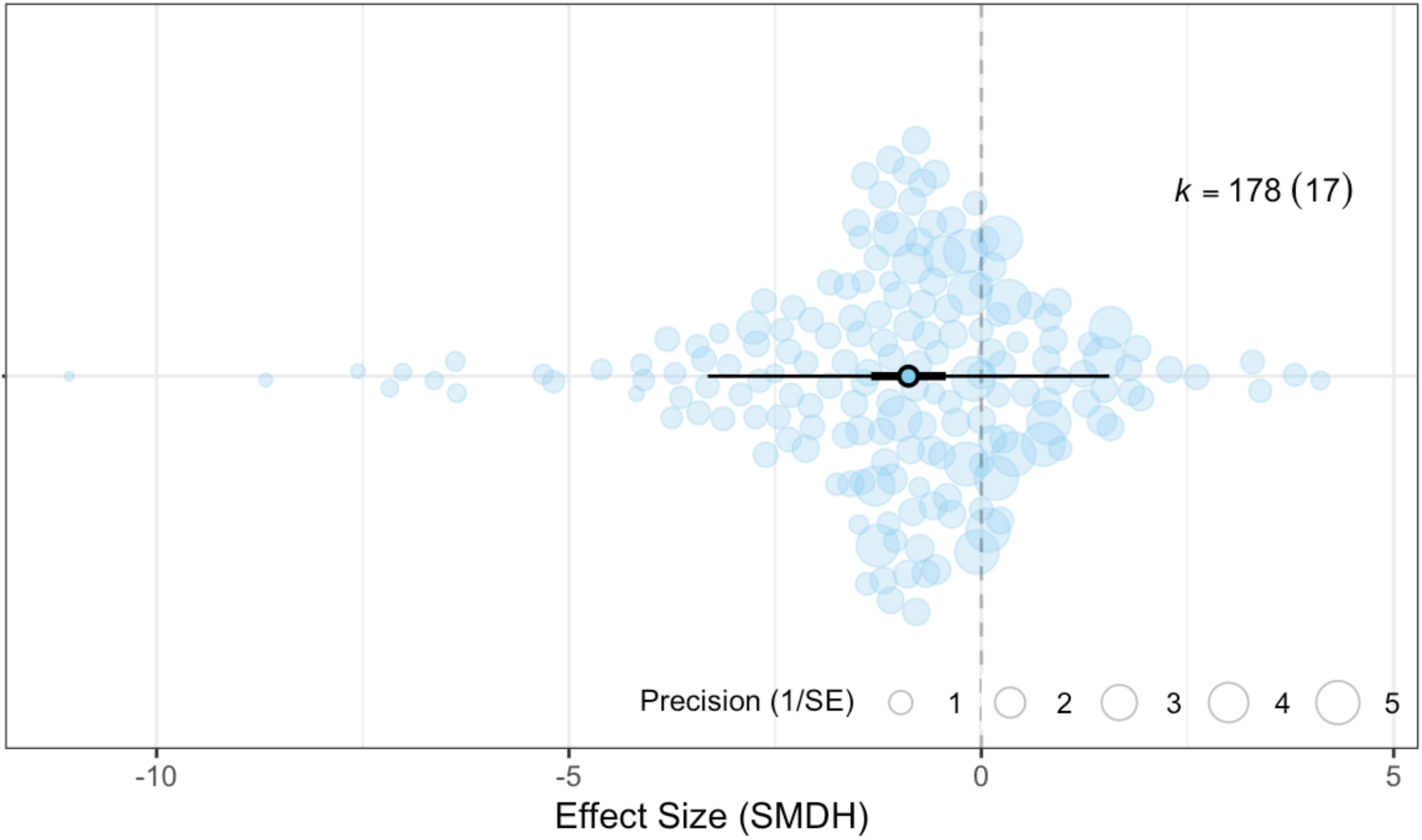
The performance of Arctic biota is negatively impacted by extreme winter events. Each data point represents one effect size. The size of each data point corresponds to the precision of the effect size, represented as 1/SE. Effect sizes (SMDH) are depicted on the x-axis, with negative values indicating negative effects of extreme events on the fitness of Arctic biota, such as increased mortality, while positive values indicate positive effects, such as higher peak flower abundance. Sample sizes are denoted as k = number of effect sizes (number of studies in brackets). Bold error bars (95% CI) signify whether the overall effect size significantly differs from zero (not overlapping zero), while light error bars depict the 95% Prediction Interval (PI) of effect sizes. The mean effect size is represented by a black circle in the centre of the horizontal line.

We compared the impact of methodological moderators to confirm that the various studies could be include in the meta-analysis. Both Bokhorst-led studies and those by other first authors showed a negative effect of extreme winter events on Arctic biota (Bokhorst: mean = −0.852, *p* < 0.010; Others: mean = −0.993, *p* < 0.050) with an insignificant difference between the two groups (*p* = 0.740). Both experimental and observational studies showed significant negative impacts of extreme winter events on Arctic biota (Experimental: mean = −0.796 *p* < 0.010; Observational: mean = −1.276, *p* < 0.010) and hence, we were able to include both study types in further analyses.

Arctic biota are negatively impacted by extreme winter warming (mean effect size = −0.655, p <0.050) and by rain-on-snow (mean = −1.476, p < 0.010, Fig. 6a), and effect sizes do not significantly differ between the two types of extreme events.

**Fig. 6.**
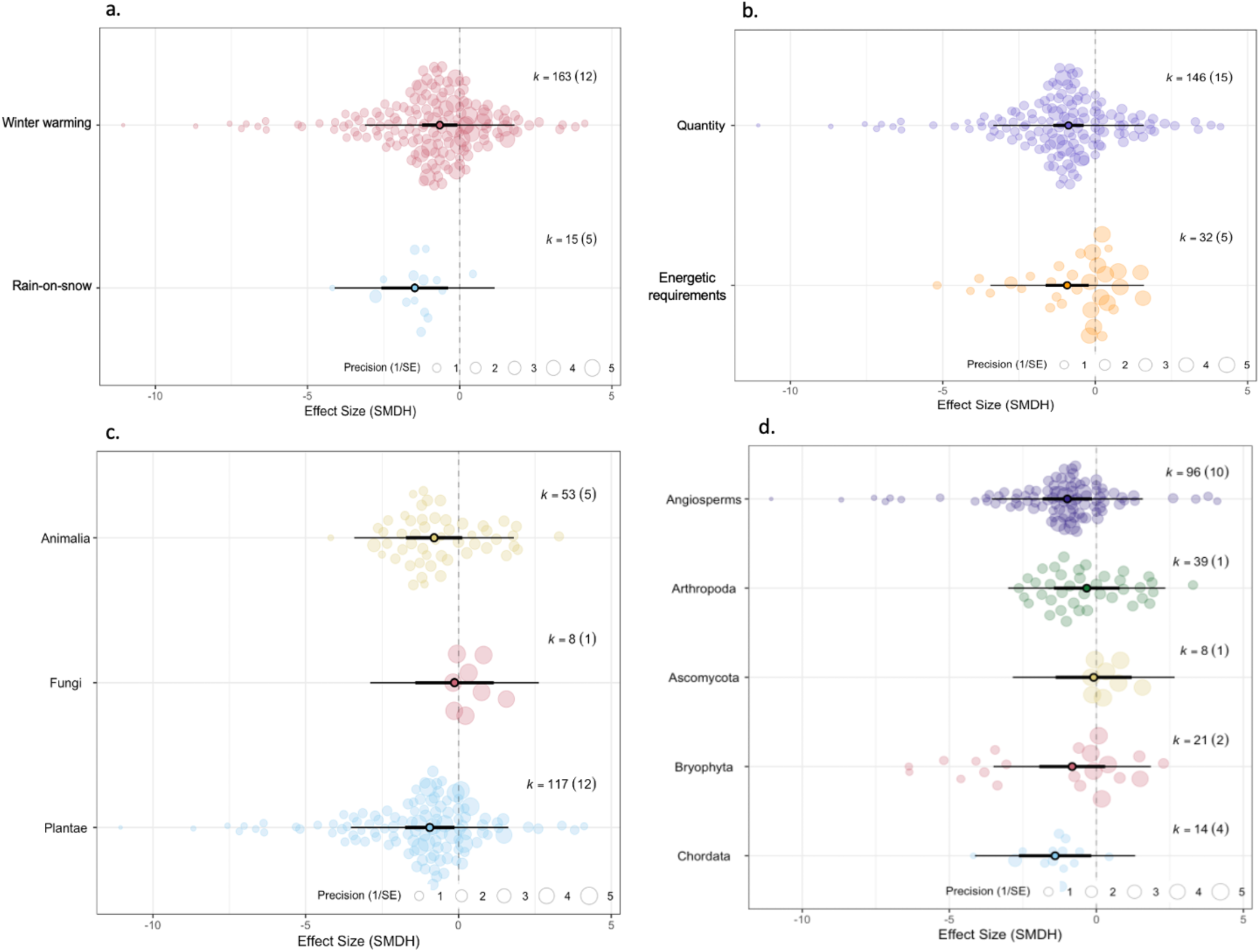
Rain-on-snow and extreme winter warming impacts on Arctic biota. (**a**) Effect sizes of Arctic biota in response to rain-on-snow and extreme winter warming on Arctic biota. (**b**) Quantitative and energy-related measures response to extreme winter events. Taxonomic kingdoms (**c**) and phyla (**d**) show varying sensitivity to extreme winter events. Each data point represents one effect size. The size of each data point corresponds to the precision of the effect size, represented as 1/SE. Effect sizes (SMDH) are depicted on the x-axis, with negative values indicating negative effects of extreme events on the fitness of Arctic biota, such as increased mortality, while positive values indicate positive effects, such as higher peak flower abundance. Sample sizes are denoted as k = number of effect sizes (number of studies in brackets). Bold error bars (95% CI) represent whether the overall effect size significantly differs from zero (not overlapping zero), while light error bars depict the 95% Prediction Interval (PI) of effect sizes. The mean effect size is represented by a black circle in the centre of the horizontal line.

Extreme winter events have equivalent impacts on quantitative responses (mean = −0.876, *p* < 0.001) and energetic requirements (mean = −0.921, *p* < 0.050; Fig. 6b). However, it is important to note that the model with ‘response variable category’ as a moderator explains the least of the heterogeneity observed in the full model (marginal R^2^ = 0.019, Q_M_(df = 2) = 13.9, *p* < 0.001; Appendix 8). This suggests that only 1.9% of the variation in effect sizes is accounted for by the response variable category. Hence, other factors are responsible for the variability in the dataset.

Taxonomic groups exhibited varying responses to extreme events, with differences observed across kingdoms and phyla. Plantae, the group with the largest sample size, was the only kingdom out of the three included that showed a statistically significant negative effect (mean = −0.942, *p* < 0.050; Fig. 6c.). At phyla level, only Chordata and Angiosperms showed a statistically significant negative response (Chordata: mean = −1.400, *p* < 0.050; Angiosperms: mean = −0.987, *p* < 0.050; Fig. Fig. 6d.) whilst Arthropoda, Ascomycota, and Bryophyta did not. The model that included phyla as a moderator explained the highest proportion of heterogeneity when compared to other models with a marginal R^2^ value of 7.23 and Q_M_(df = 5) = 10.7 (*p* = 0.059). However, a significant portion of unexplained heterogeneity remains.

## 5. Discussion

### 5.1 Trends in extreme winter climate events

Extreme climate events, whose increased frequency and intensity are driven by gradual warming trends, increasingly exceed the physiological tolerance levels of organisms leading to population declines, range shifts, and potential extinction events (Harris et al., 2018; Smale & Wernberg, 2013; Boucek & Rehage, 2014). Here, we show that extreme winter warming events have become longer and more intense whilst the trends in rain-on-snow vary across the Arctic. Whilst we report considerable variability across biotic effect sizes, we found an overall negative effect of extreme winter events on the fitness of Arctic biota. Put together, our results suggest that the shifts in the patterns of emergence in extreme winter events across the Arctic have potentially negative impacts on native biota.

The Arctic has been getting warmer with longer and more extreme winter warming events. We found that 1990-2020 experienced a significantly higher level of annual cumulative winter warming exceedance than 1950-1980. We also found that extreme warming events have been getting longer and more intense every decade since 1950. This trend is consistent with studies undertaken in the Arctic reporting that mean winter temperatures and extreme events have increased in recent decades (Callaghan *et al*., 2010; Vikhamar-Schuler *et al*., 2016; Pascual & Johansson, 2022; Rantanen *et al*., 2022). The weak decrease in annual cumulative winter warming in Eastern Eurasia is consistent with findings that extreme cold conditions have been observed in this area. Sub-seasonal Siberian cold has been linked to extreme Siberian blocking and strong East Atlantic/Western Russia patterns influencing the dipole atmospheric circulation (Luo *et al.,* 2016; Song *et al.,* 2021; Kim *et al*., 2023). The trend towards longer and more extreme winter warming events is concerning for ecological stability, as these events have strong negative effects on the fitness of Arctic biota.

At a pan-Arctic scale, the frequency of rain-on-snow events does not show statistically significant increases over the last 70 years. Regionally, trends vary considerably across the Arctic, with some regions experiencing decreasing and others increasing frequencies of rain-on-snow events. Changing precipitation patterns align with our findings of a statistically significant rise in the proportion of rain to snow in the areas of interest, supporting Bintanja & Andry’s (2017) conclusion of a broader shift towards rain-dominated precipitation across much of the Arctic. Previous studies found variable patterns in changing emergence of rain-on-snow events as a response to gradual warming trends (López-Moreno *et al*., 2021). The pattern of emergence reported herein is generally consistent with rain-on-snow events documented in the Nordic Arctic region (Vikhamar-Schuler *et al*., 2016; Bartsch *et al*., 2023), Arctic Canada (Grenfell & Putkonen, 2008), Alaska (Pan *et al*., 2018; Barstch *et al*., 2023), Siberia (Bartsch *et al*., 2010; Bartsch *et al*., 2023), and Svalbard (Vickers *et al*., 2024), amongst multiple other regions of the Arctic. It is interesting to note that the geographical distribution of studies looking at effects of rain-on-snow events on Arctic biota does not reflect their pattern of emergence. Most studies looking at rain-on-snow took place in Svalbard and Yamal, where they were found to affect reindeer populations and, in the case of Yamal, indigenous communities that rely on them (Forbes *et al*., 2016; Loe *et al*., 2016). However, none of the studies included in our meta-analysis looked at the effect of rain-on-snow in Greenland, where we detected new rain-on-snow events consistent with evidence showing unprecedented rain in Greenland (Mattingly *et al*., 2016). This mismatch is worrying, as we found rain-on-snow events to have a significant negative effect on the fitness of Arctic biota, and highlights the need for further studies in areas identified as experiencing increasing rain-on-snow.

The variability in the observed changing frequencies of rain-on-snow events across the Arctic can be explained by changing sea-ice extent, warming temperatures, and the effects of ocean and atmospheric currents (McCrystall *et al*., 2021). For example, increased moisture levels in Greenland coincide with accelerating Greenland Ice Sheet melt (Mattingly *et al*., 2016; Overland, 2022), leading to unprecedented incidents of rain-on-snow. Here we show that the west and northern coast of Northern Fennoscandia experienced a decrease in rain-on-snow, despite a significant positive trend in annual precipitation. Decreasing snow cover has been reported in Northern Norway (Rizzi *et al*., 2017), which is in line with less rain-on-snow events. Lower snowfall to rainfall ratios, shorter snow duration, and higher average temperatures have been linked to decreases in rain-on-snow frequency (López-Moreno *et al*., 2021). The Barents Sea has suffered the greatest winter sea ice retreat (Smedsrud *et al*., 2013), which has been linked to atmospheric warming and significant increases in precipitation in surrounding areas such as Yamal (Screen *et al*., 2014; Forbes *et al*., 2016). The multiple, context-dependent, interacting physical processes that can result in changes in rain-on-snow occurrence highlight the need for multi-disciplinary methods when conducting research to understand the breadth, variance, and complexity of the effects of environmental change on Arctic biota (Macias-Fauria & Post, 2018).

Limitations in our climate analysis may have led to mismatches between reported and detected extreme events and vice-versa. For example, rain-on-snow events were reported in Yamal on the 5-10 November 2006 and 8-9 November 2013 (Forbes *et al*., 2016). Whilst we identified the November 2013 event in this study, our analysis did not capture the 2006 event. This discrepancy between reported events and those identified here may be explained by some extreme events not reaching the threshold for identification used in this study. For example, on the 5-10 November 2006 in Yamal, the rainfall and snow depth thresholds for rain-on-snow were met, but the maximum temperature in the area according to the ERA5 dataset used in this study was below 1 °C, thus preventing detection of such an event in our analysis. This suggests that our detection algorithm might miss rain-on-snow events where one or more of the three variables involved in their detection (rainfall, snow depth, and air temperature) fails to cross our threshold (defined based on published literature). More generally, the relative nature of extreme climate events given their definition (that is, a value above a threshold defined from a percentile and a baseline period) make their common identification across studies highly dependent on the criteria used. This is further compounded with events defined not by one but by several variables, such as rain-on-snow. On the other hand, our climate analysis detected rain-on-snow events not reported in the literature, such as a spike in rain-on-snow in Yamal in the autumn of 2007. The mismatch between our results and the literature may be a product of biased reporting. For example, reports of rain-on-snow favour events with socio-economic consequences; Forbes *et al*. (2016) report events that impacted the livelihoods of indigenous Nenets and their semi-domesticated herds of reindeer. Hence, rain-on-snow events in areas where semi-nomadic herders and their herds weren’t present may not appear in the literature due to lack of reports. To better understand trends in rain-on-snow, future research should focus on analyses at finer geographic and temporal scales (e.g., Pall *et al*., 2019), considering key drivers of rain-on-snow (e.g., Voveris & Serreze, 2023). Additionally, enhanced monitoring efforts across the Arctic are essential for capturing the extent of rain-on-snow and their ecological impacts beyond areas of socio-economic impact (e.g., Bartsch *et al*., 2023).

### 5.2 Biotic impacts of extreme winter climate events

Our meta-analysis revealed a significant negative general effect of extreme winter events on terrestrial Arctic biota. It also revealed high heterogeneity between effect sizes, which remains unexplained. This unexplained variance supports the evidence that organismal sensitivity to extreme events is nuanced and dependent on a range of factors (Jentsch *et al*., 2009; Harris *et al*., 2018; Bokhorst *et al*., 2022). Factors like timing, severity, and duration of extreme events in relation to different life-cycle stages will likely influence an organism’s performance following an extreme event (Bjerke *et al*., 2011). None of the moderators analysed in this study succeeded to explain any substantial amount of the high variance observed in Arctic biota’s sensitivity to extreme winter events. However, it is important to note that this may be due to the limited number of effect sizes within each unique combination of moderator, which reduces the power to detect meaningful differences.

Different taxonomic groups showed varying sensitivity to extreme winter events. Bearing in mind that the effect on different taxonomic groups is sensitive to the implementation of phylogenetic corrections, and that it varies depending on the randomised bifurcation of the phylogenetic tree, we found a significant negative effect of extreme winter climate events in Angiosperms. This finding supports previous evidence that vascular plants are considerably affected by extreme events (Bokhorst *et al*., 2009; 2018). The lack of significant effects in Bryophyta and Ascomycota is also consistent with evidence that cryptograms are resistant to extreme winter events (Bokhorst *et al*., 2022). Whilst none of the studies included in our meta-analysis measured impacts of extreme winter events on bacteria, research suggests they are sensitive to extreme events (Furtak & Wolínska, 2023; Bei et al., 2023), with changes in soil microbiomes leading to significant negative effects in ecosystem productivity and future plant growth (Singh *et al*., 2020; Trivedi *et al*., 2022). The significant negative effect observed in Chordata is also consistent with studies that found that rain-on-snow events negatively impact ungulates, ptarmigan, foxes, and lemmings (Hansen *et al*., 2013; Forbes *et al*., 2016; Poirier *et al*., 2021) and have been linked to cascading ecological impacts (Stien *et al*., 2012; Layton-Matthew *et al*., 2023). Overall, our results highlight the highly complex nature of ecological responses to extreme events, which likely compounds ecological responses across ecosystems and trophic webs. Quantifying the full extent of the impact of extreme events on Arctic ecology will thus require in depth understanding of the physiological and behavioural impacts of extreme events on different species.

The paucity of data regarding the biological impacts of extreme winter events in the terrestrial Arctic is a severe limitation of our meta-analysis. Specifically, our meta-analysis only included one study investigating the impact of extreme events on lichen (Bokhorst *et al.,* 2023). Studies suggest that lichens are less susceptible to extreme events than plants, with winter warming only affecting chlorophyll fluorescence, nitrogen fixation, and photosynthetic rates in plants (Bjerke *et al*., 2011; Bokhorst *et al*., 2022). One possible explanation is the difference in growth forms and seasonal rhythms in bryophytes (Bjerke *et al*., 2011). Differing sensitivity between plant and fungi to extreme winter climate events is particularly relevant to Arctic ecology, as it may compensate for recent declines in lichen-dominated ecosystems resulting from summer warming (Walker *et al*., 2006; Rees *et al*., 2008). This hypothesis would at least partly counterbalance the effects of currently predicted climate change-enhanced increased vascular plant biomass at the expense of lichens (Cornelissen *et al*., 2001; van Wijk *et al*., 2004), as the higher resilience of lichens to winter extreme events could buffer the impacts of summer warming (Bjerke *et al.,* 2011). However, due to lack of evidence, it is impossible to conclude whether this trend is supported by our meta-analysis.

The bias within the existing literature needs to be considered in a comprehensive analysis of Arctic resilience to extreme winter events. An accurate understanding of Arctic ecology and the ways in which Arctic ecosystems will respond to climate change requires region-specific understanding. Metcalfe *et al*. (2018) found that 31% of all studies cited in terrestrial Arctic research are located within 50 km of two research stations in Toolik Lake, USA and Abisko, Sweden. They also reported that colder, sparsely vegetated sites experiencing rapid warming are under-represented within citations in the field (Metcalfe *et al.,* 2018). Similar patterns were also reported by Martin et al. in a systematic review on drivers of Arctic shrub expansion (Martin *et al.,* 2017). Uneven geographical distribution of studies was observed in our meta-analysis, where ∼65% of studies took place around Abisko, Sweden. The bias in Arctic studies has increased since the Russian invasion of Ukraine, reducing international collaborations in Russia, the largest Arctic state by extent (López-Blanco *et al*., 2024). A further likely bias in the literature stems from a focus on systems where biotic impacts of extreme winter climate events have been reported to directly affect human livelihoods. The results of our meta-analysis are only as representative as the published evidence available and likely represent a bias towards certain ecosystems and taxonomic groups. Finally, the exclusion of grey literature and of works not published in English represents a bias in our study as well. It is interesting to note that not all areas facing significant increases in either rain-on-snow or extreme winter warming are represented in our meta-analysis dataset. For example, none of the included studies discussed the effects of extreme winter events on biota in Greenland or Eastern Canada, despite these regions experiencing unprecedented climatic changes (Loeb *et al.,* 2022).

## 6. Conclusions

In this study, we synthesised the current state of literature regarding the effects of two ecologically important extreme winter events on Arctic biota and analysed geographic and temporal trends in these events. Through the combination of a meta-analysis and the analysis of climatic trends, we showed that patterns in extreme winter warming and rain-on-snow are changing considerably, and that these events have significant consequences for Arctic biota. The variability in rain-on-snow trends across the Arctic emphasises the importance of understanding key drivers such as sea ice melt and changes in atmospheric pressure and ocean currents to predict and understand future changes in precipitation. While the overall negative effect of extreme climate events on Arctic organisms is evident from the results of our meta-analysis, the high heterogeneity in our models underpins the nuanced and complex interactions between species, ecosystems, and climate. Our study also reveals substantial limitations in Arctic research, such as biases in existing literature and gaps in region and ecosystem-specific understanding that need to be addressed. Our findings reinforce the notion that the effects and emergence of extreme winter events are heterogeneous (Harris *et al.,* 2018; López-Moreno et al., 2021), however, research in these areas does not reflect this diversity. Developing our understanding of how these changing patterns will impact ecological function across the Arctic is crucial for the development of effective, region-specific conservation strategies. Future research may focus on addressing taxonomic and regional gaps to help form a more comprehensive and representative understanding of the complex dynamics driving environmental changes and their implications for the terrestrial Arctic.

