## Supplementary Appendices for "Increases in Arctic extreme climatic events are linked to negative fitness effects on the local biota"

All R scripts and data files are available online at:

https://github.com/mayalemaire/Arctic_Climate_Project

#### *Appendix 1 – The 'Arctic Region'*

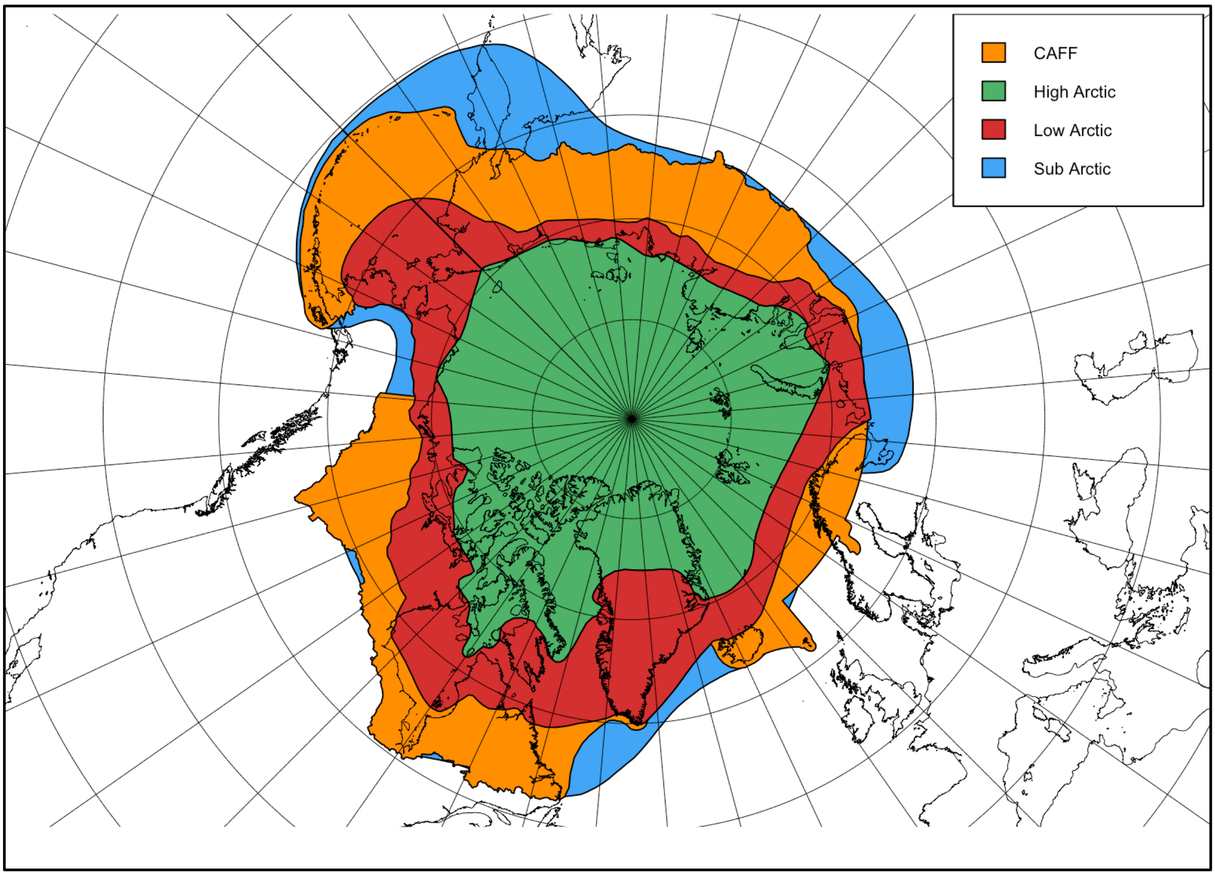


**The Arctic region** adapted from Martin *et al.*, (2022), the maximum extent of which has been used to define the Arctic in this study (i.e., the outer contour of all coloured regions). Orange = Conservation of Arctic Flora and Fauna (CAFF) boundary, Green = Arctic Biodiversity Assessment (ABA): High Arctic, Red = ABA: Low Arctic, Blue = ABA: Sub Arctic.

### *Appendix 2 – Table of most frequently used words*

| Word | Frequency | Word | Frequency | Word | Frequency |
|---|---|---|---|---|---|
| **arctic** | 1628 | studi | 876 | condit | 665 |
| **chang** | 1542 | plot | 869 | model | 633 |
| **event** | 1268 | snow | 812 | differ | 624 |
| **climat** | 1249 | **popul** | 766 | growth | 619 |
| **temperatur** | 1249 | **extrem** | 758 | **ecolog** | 609 |
| **winter** | 1175 | veget | 748 | season | 608 |
| **speci** | 1120 | high | 719 | respons | 599 |
| **warm** | 1107 | **tundra** | 713 | summer | 591 |
| year | 1094 | site | 696 | time | 573 |
| use | 990 | may | 695 | **ecosystem** | 559 |
| plant | 973 | data | 694 | caribou | 552 |
| effect | 963 | ice | 689 | observ | 527 |
| soil | 941 | area | 677 | **weather** | 525 |
| increas | 883 | fig | 671 | cover | 522 |

**The 42 most frequent words used in the 10 relevant papers identified in the scoping search to identify words for inclusion in the search string of our meta-analysis.** Pertinent words were used to produce a comprehensive search string with words in bold being selected for inclusion in the finalised search string.

***Appendix 3 – Fine-tuning and final search strings***

Fine-tuning of the search string led to the addition of “NOT (human OR energy OR infrastructure)” to each search string due to the high number of human health, infrastructure, and energy-related papers not relevant to our research questions.

The final search string for Scopus was:
TITLE-ABS-KEY (("extreme event" OR "extreme climat* event" OR "extreme weather" OR "extreme heat" OR "extreme temperature" OR "extreme precipitation" OR "rain on snow" OR "extreme winter warming") AND (tundra OR arctic) AND (ecosystem OR ecology OR species OR populations ) AND NOT (human OR energy OR infrastructure)).

The final search string for Web of Science was:
TI = (("extreme event" OR "extreme climat* event" OR "extreme weather" OR "extreme heat" OR "extreme temperature" OR "extreme precipitation" OR "rain on snow" OR "extreme winter warming") AND (tundra OR arctic)) OR AB = (("extreme event" OR "extreme climat* event" OR "extreme weather" OR "extreme heat" OR "extreme temperature" OR "extreme precipitation" OR "rain on snow" OR "extreme winter warming") AND (tundra OR arctic) AND (ecosystem OR ecology OR species OR populations) NOT (human OR energy OR infrastructure)).

***Appendix 4 - List of 17 papers included in the meta-analysis***

*Appendix 5 – Categorisation of response variables into two broader categories*

| Quantity | Energetic requirements |
|---|---|
| Abundance | Tunnel length (burrowing depth) |
| Peak flower abundance | Time to layer B (burrowing depth) |
| Alive:dead shoots ratio | Time to layer C (burrowing depth) |
| Biomass (above ground density) | Displacement (migration) |
| Shoot growth | PSII activity |
| Calf at heel (proportion of females with calf) | Photosynthetic rate |
| Winter mortality (standardised) | |
| April body mass (biomass) | |
| Berry production | |
| Survival | |
| Shoot alive | |
| Greenness | |
| Segment length | |
| Segment width | |
| Length:width ratio | |

**Response variables used in our metanalysis to determine whether rain-on-snow and extreme winter warming events in the Arctic have a significant negative effect on the fitness of native biota, categorised into two broader categories to allow for comparisons across the 17 studies examined.** Quantity-related variables assess changes in population and individual traits, while energetic requirements examine physiological and metabolic adjustments resulting from the consequences of extreme winter events.

**Appendix 6 – Arctic regions have experienced a significant increase in average maximum daily winter temperature.**

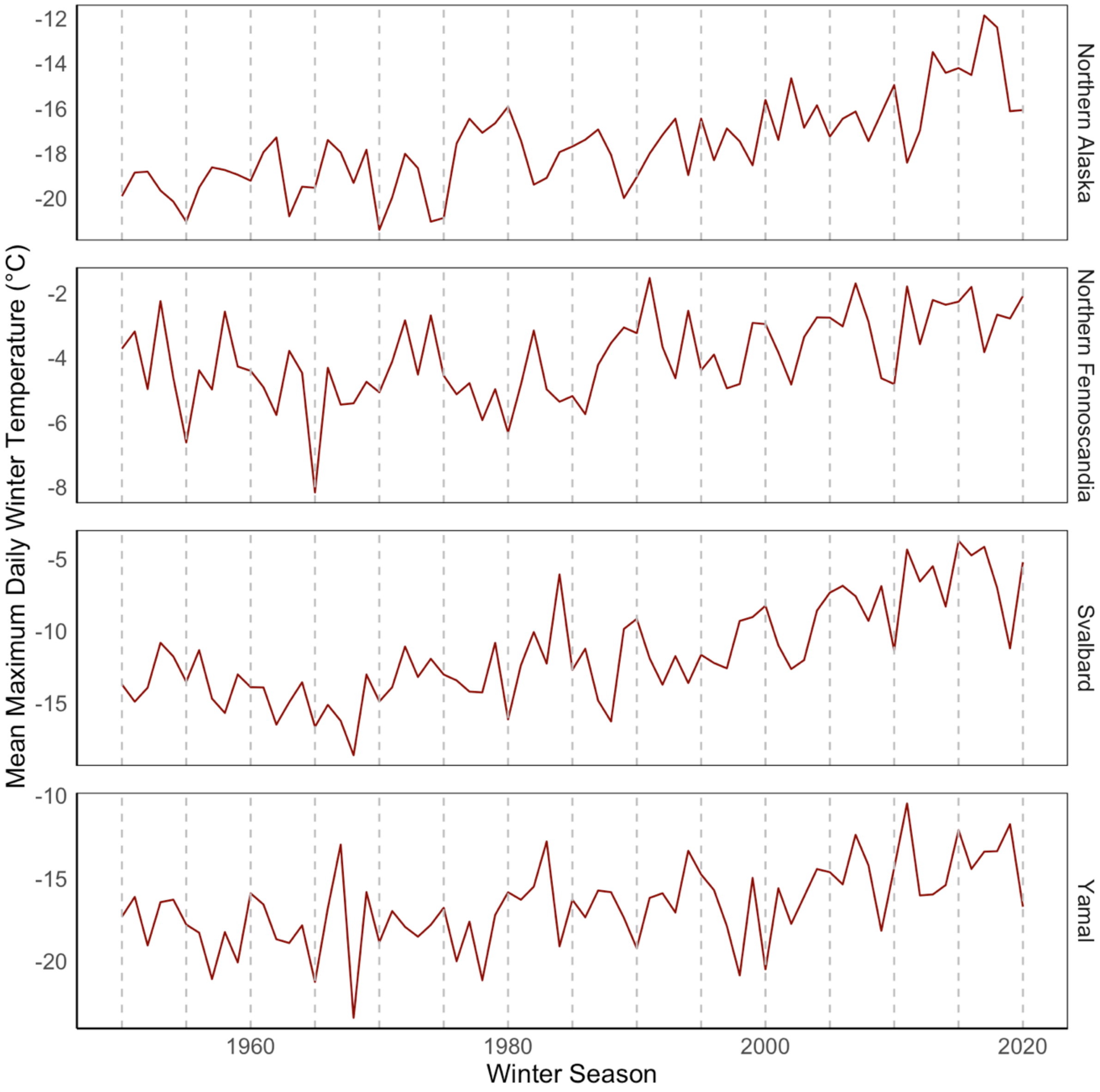


Mean maximum daily winter temperatures (°C) averaged across different Arctic areas of special interest (see Methods) across winter seasons (October to March). Grey dotted lines represent 5-year intervals. Areas of special interest are ordered alphabetically from top to bottom: Northern Alaska, Northern Fennoscandia, Svalbard, and Yamal (North-western Siberia).

***Appendix 7 – Most Arctic regions have experienced an increase in annual precipitation since 1940***

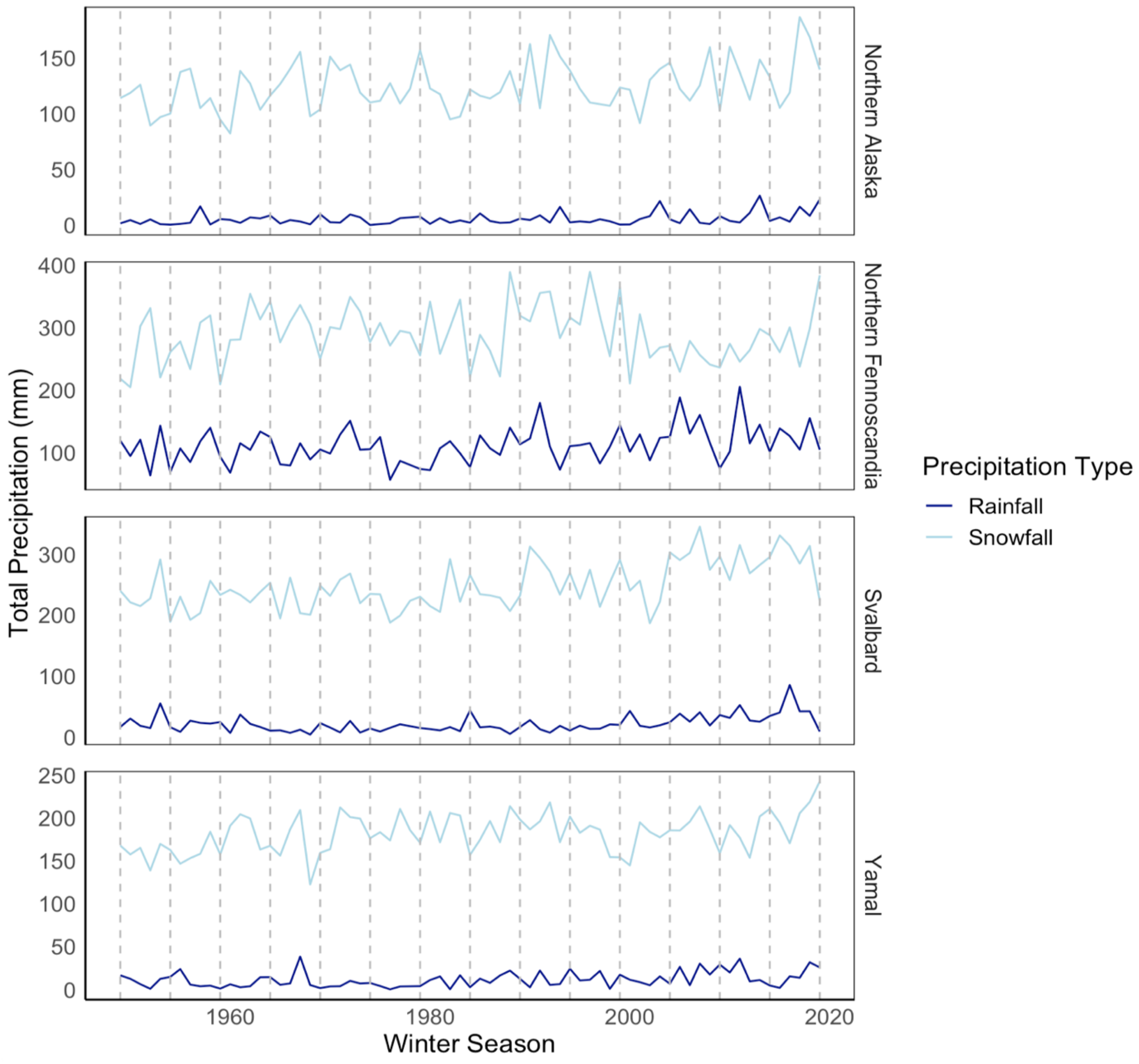


Total snowfall in millimetres water equivalent (light blue) and rainfall in millimetres (dark blue) in each winter season (October to March) for the Arctic areas of special interest (see Methods) from 1950-2020. Grey dotted lines represent 5-year intervals. Regions are ordered alphabetically from top to bottom: Northern Alaska, Northern Fennoscandia, Svalbard, and Yamal (North-western Siberia).

***Appendix 8 – Heterogeneity explained in the models with different moderators***

| **Model** | $R^2 (\%)$ |
|---|---|
| Full | 6.66 |
| Experimental Y/N | 1.27 |
| Event Type | 3.41 |
| Kingdom | 1.87 |
| Phylum | 7.23 |
| Taxonomic class | 6.76 |
| Broad Category | 0.02 |
